# Viral infection impacts the 3D subcellular structure of the abundant marine diatom *Guinardia delicatula*

**DOI:** 10.1101/2022.09.06.505714

**Authors:** Marie Walde, Cyprien Camplong, Colomban de Vargas, Anne-Claire Baudoux, Nathalie Simon

## Abstract

Viruses are key players in marine ecosystems where they infect abundant marine microbes. RNA viruses are emerging as key members of the marine virosphere. They have recently been identified as a potential source of mortality in diatoms, a group of microalgae that accounts for roughly 40% of the primary production in the ocean. Despite their likely importance, their impacts on host populations and ecosystems remain difficult to assess.

In this study, we introduce an innovative tool approach that combines automated 3D confocal microscopy with quantitative image analysis and physiological measurements to expand our understanding of viral infection. We followed different stages of infection of the bloom-forming diatom *Guinardia delicatula* by the RNA virus GdelRNAV-04 until the complete lysis of the host. From 20h after infection, we observed quantifiable changes in subcellular host morphology and biomass. Our microscopy monitoring also showed that viral infection of *G. delicatula* induced the formation of auxospores as a probable defense strategy against viruses. Our method enables the detection of discriminative morphological features on the subcellular scale and at high throughput for comparing populations, making it a promising approach for the detection quantification of viral infections in the field in the future.

## Introduction

Diatoms are some of the most successful aquatic organisms. They distribute globally in fresh and marine waters from the tropics to the poles and exhibit both benthic and planktonic lifestyles (Mann and Droop, 1996; Julius and Theriot, 2010). Diatoms comprise almost 10% of total biomass in the ocean (Leblanc et al., 2012; Bar-On and Milo, 2019) and are responsible for nearly 40% of total primary production in the global ocean, thus generating a large part of the organic carbon that sustains marine food-webs (Armbrust, 2009; Tréguer et al., 2018). Marine diatoms are also thought to contribute to ca. 40% of carbon export from the euphotic zone to depth, as part of the “biological carbon pump”, which directly influences the climate of the planet (Tréguer et al., 2018). The quantity and quality of carbon exported to depth vary according to diatom size, morphology, elemental composition, life strategy and mortality processes. Given the prominence and global impact of these unique organisms, identifying, and quantifying the mechanisms by which diatom populations are regulated is important to understand their environmental impact. Besides abiotic parameters that influence cell growth, diatom populations are also subject to the control exerted by predators (e.g., grazers) or pathogens (e.g., eukaryotes, fungi, viruses).

Viruses are the most abundant microbial forms in the ocean, and they have globally significant evolutionary and biogeochemical impacts via the infection and mortality of their microbial hosts (Fuhrman, 1999; Suttle, 2005, 200; Rohwer and Thurber, 2009). **Viruses of diatoms** were discovered rather recently (Nagasaki et al., 2004) with the isolation of *Rhizosolenia setigera* RNA virus (RsRNAV, Picornavirales, Marnaviridae). This virus possesses a tiny untailed capsid (32 nm in diameter) that enclosed a small genome made of ssRNA (9 kb), and it replicates in its host cytoplasm. Since then, viruses with similar features have been reported for 12 other diatom species (Arsenieff et al., 2022) and environmental studies suggest widespread distribution and activity of closely related ssRNA viruses in the ocean (Gustavsen et al., 2014; Miranda et al., 2016; Kranzler et al., 2019, 2021; Vlok et al., 2019). Despite their likely importance, we have only a limited understanding of the extent to which RNA viruses infect natural diatom populations and cascading impacts on diatom driven processes, mostly due to the lack of routine methods to detect and quantify infections inside cells.

In this study, we focused on the key marine diatom species *Guinardia delicatula*. It is distributed widely from the poles to the equator, mainly in coastal areas (Guiry and Guiry, 2018) and in Europe, especially along the Atlantic coast of the English Channel. In this latter ecosystem, *G. delicatula* is reported as one of the most abundant diatom species (Guilloux et al., 2013; Hernández-Fariñas et al., 2014; Caracciolo et al., 2022) and it forms massive blooms (up to 170’000 cells per litre of seawater) in spring-summer and fall, whose dynamics are changing with climate warming (Schlüter et al., 2012). Eukaryotic parasites, such as *Cryothecomonas aestivalis, Pirsonia guinardiae* (Tillmann et al., 1999), or *Aplanochytrium sp*. (Arsenieff, 2018), have been identified as potential primary agents that control these dynamics of *G. delicatula* populations in the field. (Tillmann et al., 1999; Peacock et al., 2014)A detailed study on Cryothecomonas aestivalis reported a negative relationship between the magnitude of *G. delicatula* blooms and parasite infection rate, suggesting that *C. aestivalis* play a major role in controlling blooms on the New England Shelf (Peacock et al., 2014). More recently, viral infection was identified as a potential source of mortality in *G. delicatula* populations. Several strains of *G. delicatula* viruses belonging to the order Picornavirales (genus Bacillarnavirus) have been isolated in coastal waters off Roscoff. Like many other diatom viruses, these pathogens only infect a limited number of strains within their host species (Tomaru et al., 2015). During their infection cycle, Guinardia viruses **replicate in their hosts’ cytoplasm** and are released in a timely manner (<12 h after infection) with complete host lysis usually detected within 3 days post-infection in the laboratory.

Although they are believed to play an important role in regulating *Guinardia* populations (Arsenieff et al., 2019), there are to date no existing methods to assess the extent of viral diatom infections with single-cell resolution. Here, we used automated high content 3D imaging to identify subcellular morphological changes during viral infection in order to expand our understanding of diatom responses to infection. We set up an infection experiment and focused specifically on changes in cell morphology (nucleus, chloroplasts) in infected cultures compared to controls using multicolour confocal microscopy and quantitative image analysis. This imaging method permits optical sectioning of samples and is thus particularly well suited to visualise fine structures in 3D within cells (White et al., 1987, 199). We combined confocal microscopy with automation of acquisition, which has been shown to provide a good balance between throughput and resolution for marine planktonic species in the 5-200 μm size range (Colin et al., 2017). From the image data, we derived morphometric analysis of organelle volumes and cell degradation. Microscopy monitoring of hundreds of individual cells was combined with measurements of viral growth parameters (virus and host cell concentrations) and global cell physiology (photosynthetic efficiency) during the infection cycle. Our approach enables the quantification of discriminative features on the subcellular scale for comparing populations. The morphology and biomass of the host population underwent measurable changes. We furthermore observe the production of auxospores as a potential virus escape mechanism.

## Materials and methods

### Algal and viral culture

The marine diatom strain *Guinardia delicatula* was obtained from the sea pier (Estacade) of Roscoff and maintained in the Roscoff Culture Collection (RCC3083). *G. delicatula* was cultured in K+Si medium (Keller et al., 1987). The culture was maintained at 18°C under a 12h light / 12h dark cycle at 100 μmol photons/m^2^s.

The clonal RNA virus GdelRNAV-04 (RCC5812) lytic to *G. delicatula* was used for this study. The viral isolate was transferred every week by inoculating an aliquot (10% v/v) of the viral suspension into an exponentially growing host suspension. The mixture was incubated under the host culture conditions (see above) until complete host lysis.

Algal growth was monitored every day by quantifying the host natural fluorescence *(in vivo* Chlorophyll a) using a spectrofluorometer equipped with a multiwell plate reader (Spark-Cyto, TECAN, Switzerland). After 10 days, wells with fluorescence lower than 50 % of that of the control were considered as lysed.

### Measurement of infection kinetics

400 mL culture of *G. delicatula* in exponential growth was established and divided into two equal aliquots of 200 mL. One aliquot was infected with 20 mL of freshly produced viral lysate. The second served as a control that was amended with 20 mL of K+Si medium. Both cultures were maintained at 18°C in sterile conditions under a 12h light / 12h dark cycle at 100 μmol photons m^-2^ s^-1^. Samples for viral titer, F_v_/F_m_, and microscopy analysis were taken twice daily (10:00 and 16:00) for 5 days (Supp. Fig. S1).

### Viral titer

The viral titer (i.e. concentration of infectious viral particles) was measured using the most probable number (MPN) assay (Reed and Muench, 1938): The viral titer was obtained by dilution to extinction of the virus sample in an exponentially growing host suspension in a 48-multiwell plate. Therefore, a 50 μL sample was inoculated in 450 μL of the host culture, mixed and serially diluted in 10-fold increments (from 10^1^ to 10^14^). Dilution series were made in triplicate. The Most Probable Number was calculated using online software (*mostprobablenumbercalculator.epa.gov*).

### Photosynthetic efficiency

The quantum yield of photosystem II of algal cells (F_v_/F_m_) measures the proportion of open photosystems II in a photosynthetic sample and is used to evaluate the activity of the light phase of the oxygenic photosynthetic process. It is thus used as a proxy for the health of the photosynthetic machinery. This parameter was determined using a pulse amplitude modulated fluorometer (Phyto-PAM, Walz, Effeltrich, Germany) connected to a Chart recorder (Labpro, Vernier). 2 mL samples were collected in plastic cuvettes and stored in the dark during a minimal relaxation time of 5 min. After this incubation time, samples were placed in the fluorometer chamber, and the non-actinic modulated light (450 nm) was turned on in order to measure the fluorescence basal level, F_0_. A saturating red-light pulse (655 nm, 4000 μmol quata /(m^2^s), 400 ms) was applied to determine the maximum fluorescence level, F_m_. The maximum quantum yield, F_v_/F_m_, was determined as

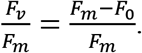

### Cell mounting and fluorescent labelling for microscopy

For each time point, 2 mL of the infected and control cultures were fixed in paraformaldehyde (Electron Microscopy Sciences, ref. 15714, 1% final conc.) and glutaraldehyde (Sigma, ref. G5882-100ML, 0.25% final conc.). After an incubation of 15 min at 4°C, 100 μl of sample solution was loaded in chambered coverglasses with 8 wells (Nunc LabTek II or ibidi μSlides), which had been pre-coated with poly-L-lysine to facilitate cell attachment. The fixed sample solution was diluted with 750 μl filtered seawater (FSW) and slowly centrifuged for 2 min to sediment all cells. Two fluorescent dyes were used: Hoechst33342 to label DNA (Thermo Scientific Pierce, Excitation: 405 nm and Emission: 415-475 nm) and StrandBrite Green to label single-stranded RNA (AAT Bioquest, Excitation: 488 nm and Emission: 515-545 nm). In each well 50 μL of FSW containing 10 μM Hoechst33342 and 1.5 μL StrandBrite Green were added. (An overview of the bioimaging pipeline is shown in Supp. Fig. S2.)

### Tests of RNA-specific fluorescent dyes

A particular challenge in detecting RNA virus replication hotspots inside photosynthetic diatoms by fluorescence microscopy is the fluorescent background: the bright and spectrally broad background autofluorescence of their chloroplasts and an omnipresent background of low-concentrated RNA inside cells. A suitable ssRNA marker and careful adjustment of spectral settings was thus a crucial prerequisite for this study. We tested a series of nucleic acid dyes with very similar spectral properties (SybrGreen, PicoGreen, StrandBrite and SYTO RNASelect) with regard to (i) their compatibility with formaldehyde fixation and (ii) their specificity for ssRNA inside infected and non-infected diatoms in comparison to a DNA stain (Hoechst33342). Further details are found in Supp. Mat. 6. We achieved the best signal-to-noise ratio with StrandBrite. Hence, this fluorescent dye was chosen for all subsequent experiments.

### Abundance counts

Cell counts were extracted from low-resolution overview images (10x objective, Supp. Fig. S3) of known volumes from each sample by automatic particle counting in FIJI (Schindelin et al., 2012). A detailed description of the steps can be found here: *imagej.net/imaging/particle-analysis*. In brief, after light Gaussian blurring (σ=0.5), cells were segmented from the background by thresholding based on their Chlorophyll autofluorescence (Otsu dark method). The resulting binary masks were dilated and any holes in detected objects were filled. At low resolution, fewer subcellular features are available for the classification of “healthy” or “dead” cells. Thus, all cells with bright chloroplasts and within typical shape parameters for “healthy”*G. delicatula* (size>30.00; circularity=0 to 0.6) were counted.

Bacteria concentrations were manually counted from the Hoechst33342 fluorescence channel of high-resolution images. Their concentrations were then calculated by extrapolation from the mean counts of all imaged areas to the total volumes loaded into each imaging chamber.

### Automated image acquisition

A confocal microscope (Leica SP8, Germany) equipped with 63x oil NA 1.4 objective was used. For high-content screening of many individual cells, a dedicated experimental “Mark&Find” template was designed and pre-tested with the Leica MatrixScanner automation software. The imaging experiment consists of three essential steps:

1. Acquisition of a reference focus map,
2. a fast low-resolution scan to detect cell positions, and
3. a 3D high-resolution scan at selected positions only.

During the high-resolution scan, cell morphologies post-infection automatically according to the following predefined settings: 68 z-planes were recorded over a total height distance of 20 μm for each cell. At each of these z-planes, fluorescence was read out in 3 fluorescent channels and one transmission light channel:

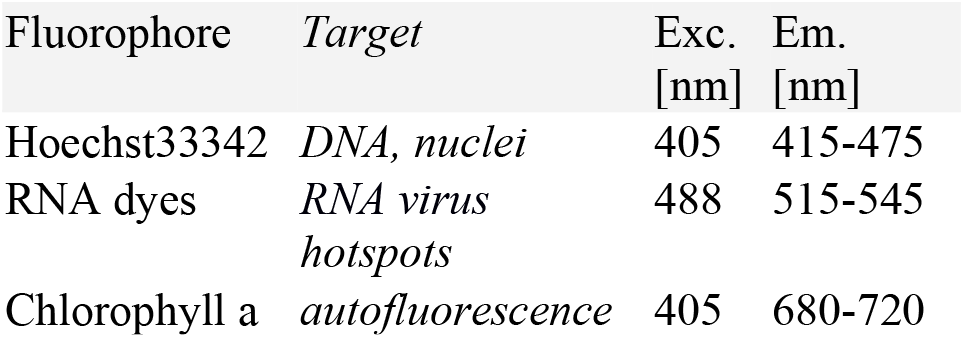

To minimise fluorescence crosstalk between recorded channels, A two-step sequential acquisition was designed: First RNA dye and transmitted light channel were recorded simultaneously, then Hoechst33342 and Chlorophyll autofluorescence. Per time point, around 50 cells from infected samples per time point were chosen for high-resolution 3D imaging and around 20 cells from control samples, thus more than 550 individual cells in total during the course of a viral infection.

### Image processing for visualisation

For 2D visualisation of multi-colour 3D datasets from confocal microscopy, datasets were batch-processed in FIJI (Schindelin et al., 2012). The full annotated script can be found on *githubxom/mariesoopy/GuinardiaGdrallnfection*. In brief, for each z-stack recorded, a 2D maximum projection was calculated for each fluorescence channel and a minimum projection for the transmission channel. For an overview collection of individual cells (Fig. 2 & Supp. Fig. 2), the dominant cell orientation in each image was detected by Fourier component analysis (Tinevez et al., 2018), and cells were aligned along the vertical axis and cropped to their bounding box. In case of ambiguities in the image, we rotated them by hand.

### 3D morphometry analysis

The total volume of *G. delicatula* cells *V_Gui_* can be approximated by their cylindrical shape (Leblanc et al., 2012), which is described by two parameters, its diameter *d_c_* and height hc [μm] (see also Supp. Fig. 4), both of which were measured from image data:

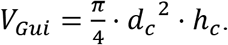

Only the volumes of intact frustules which still contained biomass were considered.

To compare the volume of entire cells to subcellular morphological changes, we also measured the volume of nuclei and chloroplasts in single cells directly from 3D image renderings:

The biovolume of nuclei was estimated from voxels (3D pixels) showing Hoechst33342 fluorescence signal. If several such volumes were detected inside one cell (i.e., in the case of bacterial invasions) only the largest detected volume was considered. The volume of chloroplasts was estimated from the sum of voxels showing chlorophyll autofluorescence. As a metric for the shape of chloroplasts, we analysed the sphericity of their 3D renderings. It is a measure of how closely their shape resembles that of a perfect sphere. Sphericity *Ψ* is defined as the ratio of the surface area of a sphere (with the same volume *V_O_* as the given object) to the actual surface area *A_O_* of this object:

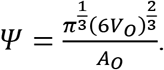

A sphere has a *Ψ*=1, while elongated objects have lower values between 0 and 1. Usually, several chloroplasts were detected inside the same cell. We then took their mean sphericity. 3D renderings and quantifications of organelle morphologies were conducted with custom macros in FIJI based on the 3D ImageSuite plugin (Ollion et al., 2013) and cross-checked with Imaris 3D image visualization and analysis software (Oxford Instruments, UK). For each time-point, we tested for significant statistical differences (t-tests) between the observed morphologies infected and control population. All statistical analysis was conducted in RStudio.

## Results

### Diatom abundances and viral titer

In the uninfected control samples, the total abundance of cells (any elongated *Guinardia delicatula* containing plastids) increased from 0 to 92 hours post-infection (hpi) until it reached a maximum of 29.6×10^3^ cells per mL (Fig. 1A). In the infected samples, cell abundance dropped from 14×10^3^ (26 hpi) to a minimum of 2.6×10^3^cells per mL (92 hpi). (And most of these remaining *G. delicatula* cells showed severe deformations upon closer inspection, see also Fig. 2).

**Figure 1.**
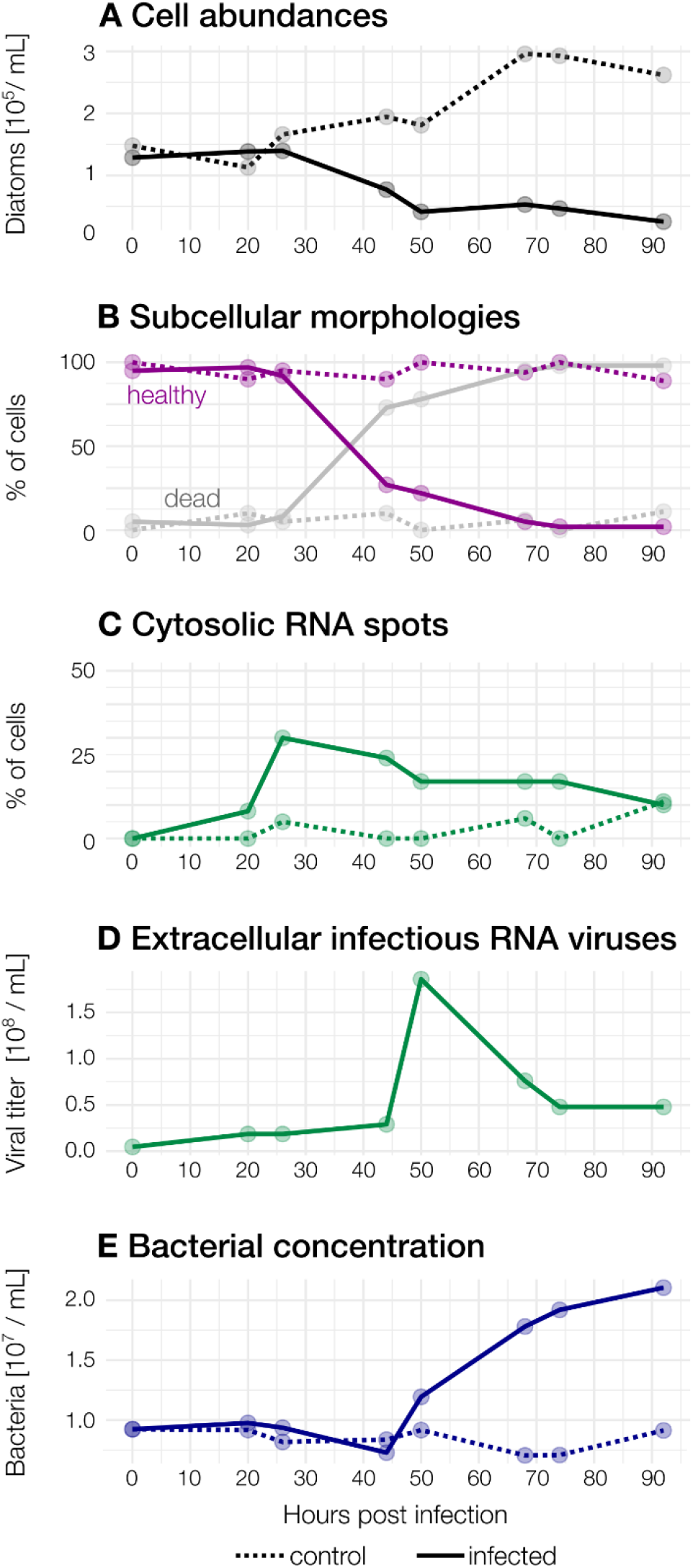
Observed abundances during the course of the infection. *Dashed: control, solid: infected* **(A)** Diatom abundances in infected and control samples. (**B**) Subcellular morphologies. In infected samples, the proportion of “healthy” diatoms declines rapidly after 26hpi, but not in control samples. Cells were counted as “dead” when they showed a broken frustule, no nucleus and/or were heavily colonised by bacteria. **(C)** Cytosolic RNA spots. (**D**) Concentration of infectious extracellular RNA viruses. (**E**) Bacterial concentrations increase from 50 hpi in the infected culture.

**Figure 2.**
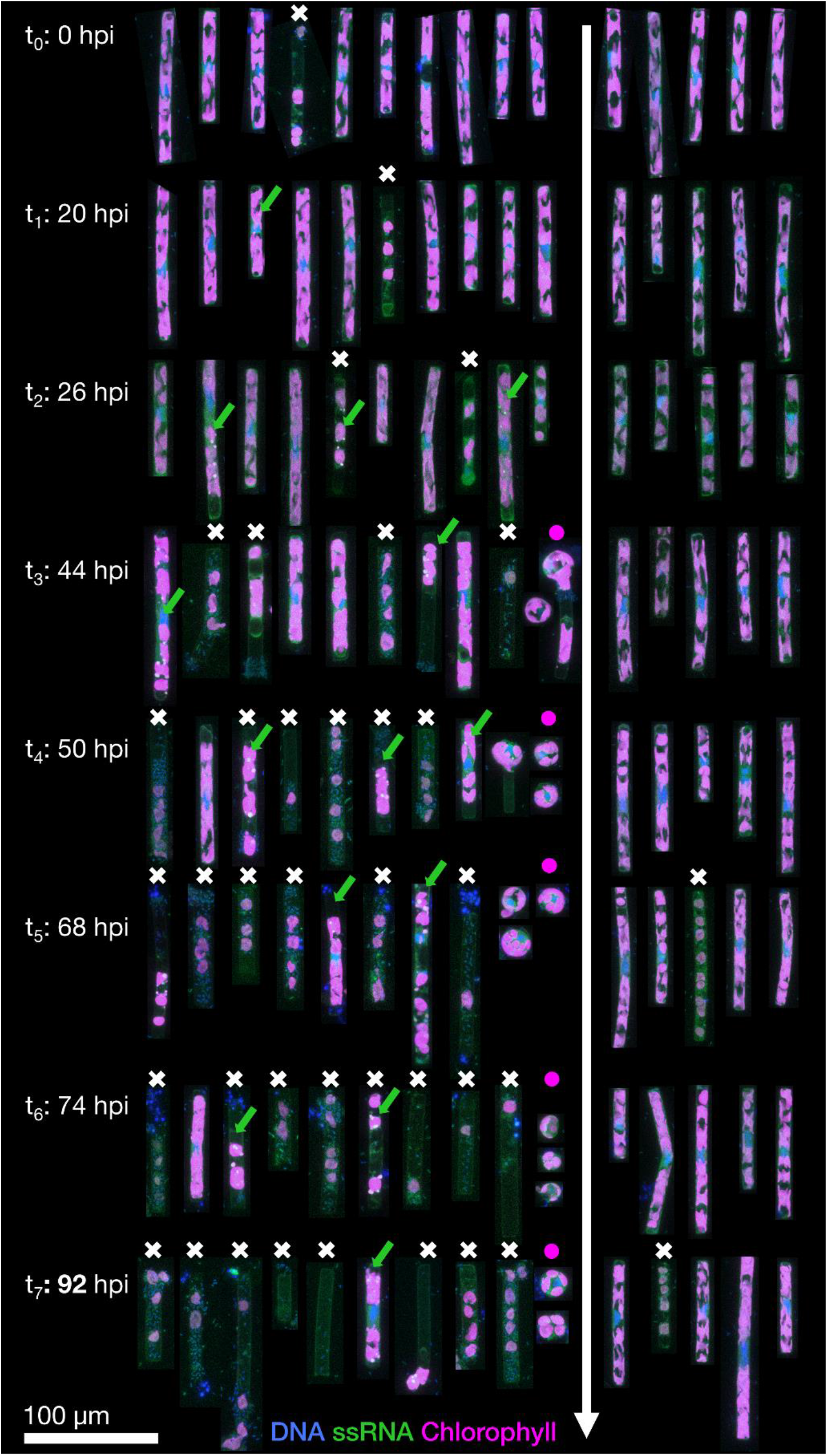
Representative Guinardia delicatula cells from samples infected with GdelRNAV-04 (left) and a non-infected control culture (right). Each cell is the depicted as the maximum projection of a corresponding 3D confocal dataset, with blue showing DNA and nuclei, green single-stranded RNA (including large viral accumulations, *green arrows*) and magenta autofluorescence from chlorophyll. Dead cells are highlighted by a *white cross*, and the increased formation of auxospores by *purple dot*.

Viral concentrations slightly increased (from 4.59×10^6^ to 1.86×10^7^ mL^-1^) within 20 hpi compared to t_0_ (Fig. 1B) concomitant with *Guinardia* cell abundances and without detection of cell lysis.

The maximal viral concentration was observed at 50 hpi with a concentration of 1.86×10^8^ viruses per mL, indicating the extracellular release of virus progeny. After 50 hours, a rapid decline of virions was observed (7.6×10^7^ per mL, 68 hpi). At the end of the experiment, the concentration of extracellular viruses stabilised at 4.79×10^7^ per mL.

### Morphological and physiological changes

From the 3D image data of each cell, we monitored morphological changes in control and infected cultures (Fig. 2) and quantified the deformation of their subcellular components throughout the viral infection (Fig. 3A-C). Complementary physiological measurements of F_v_/F_m_ were taken in parallel to relate them to observed morphological changes.

**Figure 3.**
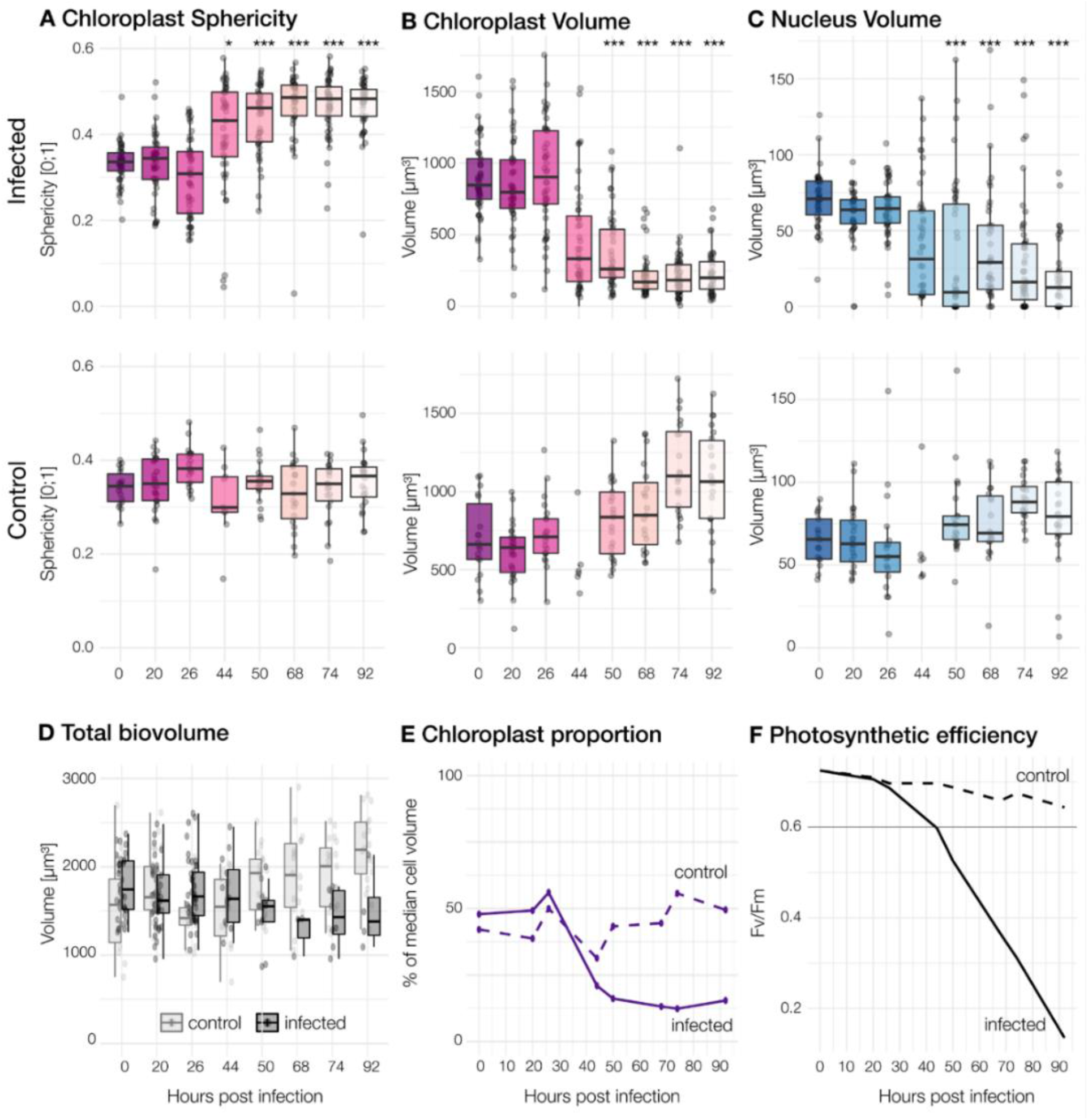
Morphological and physiological changes for different timepoints. Each dot represents one cell. *Top row:* Infected cells, *Second row:* Control cultures. Significant differences between infected and control population are marked by *** for *p* < 0.001 < and * for *p* < 0.05. **(A)** Sphericity of chloroplasts **(B)** The summed volume of chloroplasts in each cell. (C) Volume of nuclei. (A slight defocus in the control at 44h problem led to fewer quantifiable 3D datasets of individual cells.) **(D)** The total biovolume per cell, estimated from intact frustule size. **(E)** Relative proportion of chloroplast volumes compared to total volume per cell. **(F)** Maximum quantum efficiency of photosystem II (F_v_/F_m_; values under 0.6 indicate stress)

Cells in all control samples had irregularly lobed chloroplasts pressed against the cell wall and a single nucleus (stained by the DNA dye Hoechst33342) located at the centre of the cell (Fig. 2) all along the experiment. Cells with these characteristics were further considered “healthy”. Chloroplast median sphericity of these cells remained between 0.3 and 0.4 throughout the experiment while the median volume of chloroplasts per cell was between 600 and 1100 μm^3^ with the highest values recorded at the end of the experiment (92 hpi). The median nucleus size was about 60 μm^3^ in the experiment’s early stages and slightly increased afterwards. Some rare cases of dead cells were observed in control cultures.

In **infected samples**, during the early stages of infection (0 to 26 hpi), cell morphology was highly similar to that observed for non-infected cultures although median chloroplasts volume per cell was slightly higher (above 700 μm3) (Figs. 2 & 3A-C). Morphological changes were observed in a few cells at 26 hpi and then in an increasing number of cells until the end of the experiment in infected cultures:

First of all, chloroplasts that were initially irregularly lobed (as in control cultures) became more rounded and less pressed against the cell wall (Fig. 4). Median **chloroplast sphericity** increased at 44 hpi to 0.45 and stabilised around 0.49 from 68 hpi. From 50 hpi, the observed morphological changes become significantly different between control and infected samples (Fig. 3).

**Figure 4.**
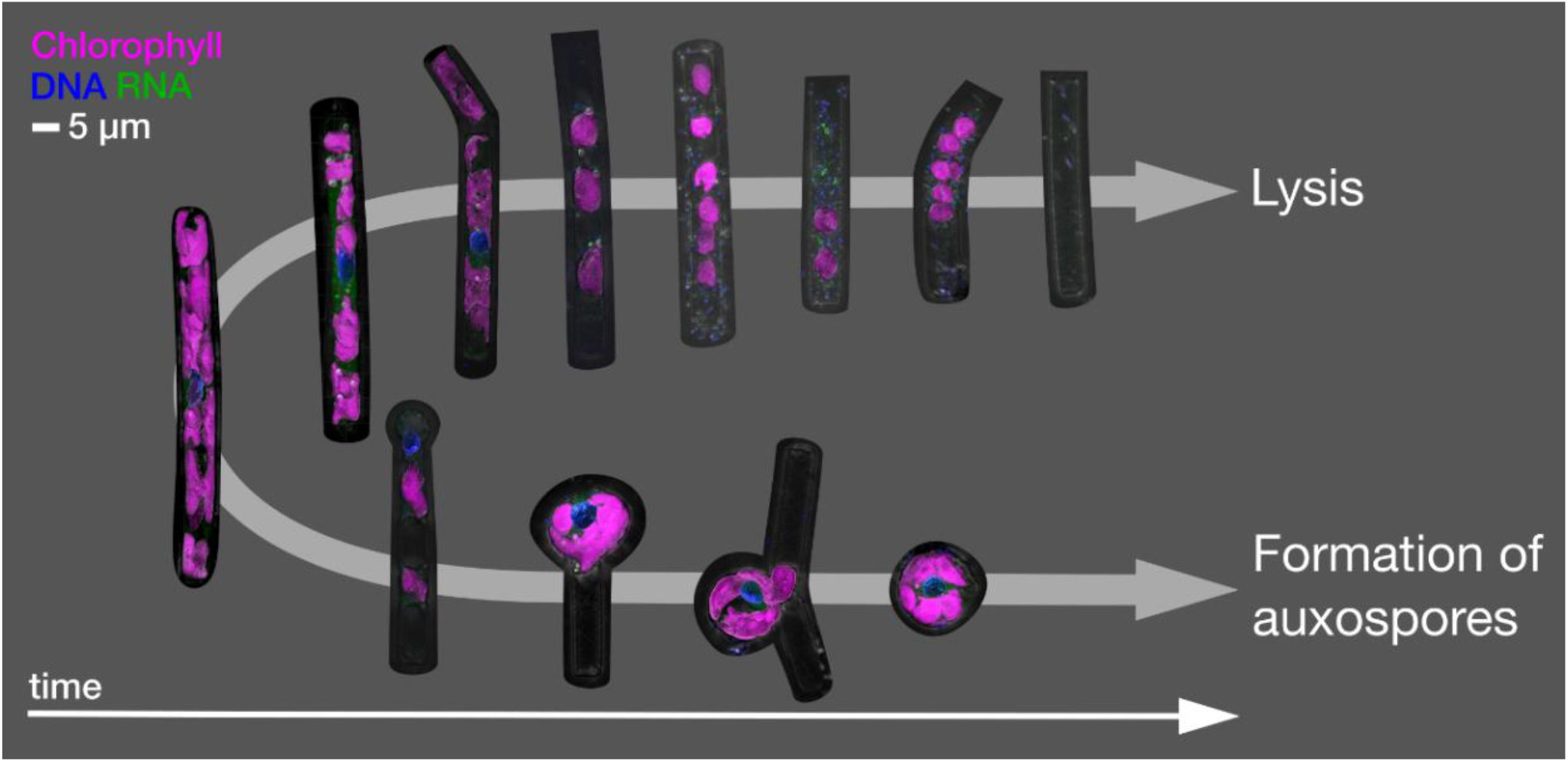
Main cellular trajectories in the infected culture. *Upper arrow:* typical morphologies of virally lysed cells. *Lower arrow:* auxospore formation. Shown are 3D renderings of selected representative morphologies.

Second, distinct **cytosolic RNA spots** (labelled by RNA dye StrandBrite) were observed from 26 hpi in cells that appeared healthy (Fig. 2). At 26 hpi, RNA spots were observed in 30% of the cells from infected samples (Figs. 1 & 2). Around a quarter (24%) of cells in infected culture showed RNA spots at 44 hpi. At 50 hpi, cytosolic RNA spots were still detected in nearly a fifth (17%) of cells, regardless if they appeared healthy or not. At this stage, bacteria frequently appeared in the vicinity of infected diatom cells and inside broken frustules, possibly contributing to the deformation of infected diatoms.

Third, at 50 hpi, we also observed an additional type of circular cells, likely **diatom auxospores**, characterised by a diameter of 15 and 25 μm, lobed chloroplasts and a nucleus located at the centre (Fig. 2). We did not detect RNA spots inside these auxospores. From 68 hpi until the end of the experiment (92 hpi) most *G. delicatula* cells were dead or severely damaged and colonised by bacteria in the infected culture (while few dead cells were detected in the control culture (Fig. 1).

The **photosynthetic machinery** in infected cultures maintained an optimal F_v_/F_m_ value for 26 hours post-infection. From 26 to 96 hpi (end of the experiment), this ratio progressively decreased from 0.69 to 0.14 (Fig. 3F). Thecontrol culture had an optimal F_v_/F_m_ throughout the experiment, varying from 0.73 to 0.64.

### Cell mortality over time

We observe a decline of diatom abundance (Fig. 1A) in the infected samples after 26 hpi. The relative proportion of healthy cells, dead cells (characterized by an absence of a nucleus, a broken frustule, and the invasion of bacteria) and cells containing RNA spots were estimated from 2D projections of high-resolution 3D images of infected and non-infected cells: In infected samples, 95% of the cells were healthy at t_0_ (Figs. 1B & 2) and no cytosolic RNA spots were detected (Fig. 1C). Between 26 to 55 hpi, the proportion of healthy cells dropped from over 92% to 27%. At 44 hpi, dead cells largely dominated in the infected samples and reached 98% at 92 hpi, concurrent with the sharp increase in viral titer in the extracellular medium between 42 and 50 hpi (Fig. 1D). Increasingly only empty frustules were observed. The percentage of cells with cytosolic RNA spots reached a maximum at 26 hpi (30%). It decreased slowly afterwards and reached 10% at the end of the experiment. For the control sample, the proportion of healthy cells was above 80% during all the experiments (Fig. 1B). Bacteria were detected around and inside most dead *G. delicatula* cells, potentially interfering with the virus mediated infection process and likely feeding on the available organic matter. From 50 hpi until the end of the experiment the bacterial concentration (Fig. 1E) steadily increased.

### Biovolume declines during the infection

The median volume of diatom cells based on the cylindrical shape of intact frustules (Fig 4D) slowly decreased in infected cultures (from ~1700 to ~1400 μm^3^) while it continuously increased in control cultures (from ~1600 to ~2200 μm^3^). Our observations and measurements of subcellular organelles reveal, however, that the relative size of chloroplasts inside the frustules continuously decreased (Fig. 3E & Suppl. Table S4a): While they accounted for 39% to 56% of the volume in all control samples, we saw a drop from 48% to 15% in infected samples. The proportion of nuclei is small compared to chloroplasts, but likewise dropped from 4% to 1% during the viral infection and remained approx. 4% in control cultures. This suggests that in later stages of the infection the effective biomass per diatom at the end of the experiment might be overestimated up to three-fold in infected cells when calculated based on diatom geometry.

## Discussion

### Morphological changes linked to viral infection

We observed that infectious GdelRNAV-04 virions (virus progeny) are released extracellularly after 20h, suggesting that the latent period is shorter than 20 hours. As reported for GdelRNAV-01, virions were released prior to Guinardia host lysis during the first 20 hpi (Arsenieff et al., 2019). Such pattern of virus production prior to host cell lysis is commonly reported in diatom-virus systems (Shirai et al., 2008; Tomaru et al., 2014; Kimura and Tomaru, 2015; Arsenieff et al., 2022). Virion concentration increases progressively to reach a maximum at 50 hours. Viral infection ultimately induced complete host lysis after 68 hours of infection. Our microscopy monitoring showed **important morphological changes during infection**: *Guinardia delicatula* cells in the control samples were cylindrical and possessed irregularly lobed and strongly pigmented chloroplasts. Infected cells remained mostly intact for 26 hours. From this time, we recorded an impairment of the photosynthetic machinery (F_v_/F_m_ <0.6), while the shape of chloroplasts still remained mostly intact. These changes were concomitant to the detection of dense RNA spots in the host cytoplasm. It is likely that this accumulation of RNA in infected host cells corresponded to the formation of viral arrays which have already been observed in electron microscopy images of GdelRNAV-01 (Arsenieff et al., 2019). We observed a peak of such spots just before large amounts of cells in infected samples were lysed. It would be interesting to develop single-cell correlative microscopy (CLEM) protocols in order to get insights into the processes of subcellular viral accumulations.

The degradation of photosynthetic pigments and reduction of photosynthetic activity are typically recorded during the late stage of phytoplankton death and, thereby, cannot be used as a proxy for viral infection (Veldhuis et al., 2001). By contrast, the formation of dense RNA spots, which likely corresponds to the accumulation of mature RNA virions, stands out as an interesting feature to quantify virally infected *Guinardia*. Because most of the host population collapses at the end of the experiment, cells that exhibit dense RNA spots can represent only a fraction of virally infected cells (because mature particles are only visible at the late stages of the infection cycle). Sporadically, RNA spots were visible in non-infected cells, and it seems likely that other cellular processes could also lead to local RNA enrichments. It will be interesting to investigate them in more detail with additional analyses, such as qRT-PCR or virus-FISH (Allers et al., 2013) targeting GdelRNAV genes or by correlative electron and light microscopy.

### Interplay of bacterial and viral infections

Viral infection was shown to induce drastic cellular alterations such as modification in resource assimilation and redirection or ultrastructural rearrangements in a wide variety host cells (bacteria, animal or plant cells) (Netherton and Wileman, 2011; Mayer et al., 2019; Warwick-Dugdale et al., 2019). In line with this, the physiological and morphological changes detected during the experiment likely reflect the metabolic hijacking and host responses during viral infection. However, the observed increase in bacterial concentration (Fig. 1E) suggests that the composition of the bacterial community might have been modified and possibly contributed to the morphological alterations of diatoms during the experiment. For example, we cannot rule out that infected cells with altered metabolism (and exudate) selected for algicidal bacteria strains which in turn could induce changes in diatom morphology and ultrastructure. The relative impact of viruses versus bacteria on host morphology would certainly require additional experimentations. Yet, the chronology of our observations strongly supports the hypothesis that observed changes and lysis were mostly triggered by viral infection and that bacteria rather benefited from the release of lysis products.

### Viral infection and life cycle phase transition

Another finding of this study is the **formation of auxospores** 44 hours after the infection while only vegetative cells were observed in control cultures. Auxospores typically play a role in growth processes, sexual reproduction, or dormancy. Diatoms are diploid organisms with a monogenetic life cycle. They reproduce through prolonged periods of mitotic divisions (Mann and Droop, 1996; Kröger and Poulsen, 2008; Julius and Theriot, 2010). During mitosis, new valves are formed inside the mother cell and each daughter cell will receive its epitheca from the mother cell and a newly formed hypotheca (Mann and Droop, 1996; Julius and Theriot, 2010). As a result, the daughter cell cannot be larger than the initial cell. This phenomenon generally causes a gradual decrease in cell size. When the size decreases to a minimum, it triggers sexual reproduction through the formation of auxospores which restores the original cell size (Kröger and Poulsen, 2008).

Concurrent with the appearance of auxospores, we found significant differences in the shape of chloroplasts between infected and non-infected cells (from 44 hpi) which became more rounded and less attached to the cell wall. This could indicate the mass onset of sexual reproduction in the infected diatom population (Crawford, 1995). Such induction of sexual reproduction by viral infection has been reported recently in another diatom-virus system (Pelusi et al., 2021). In the host-virus system, both partners are constantly changing and adapting in response to each other’s attack and defence strategies. This mutual selection refers to the Red Queen hypothesis, or the dynamics of the arms race and especially “sex against parasites”. In this hypothesis, hosts have developed sexual reproduction to maintain genetic diversity against rapidly evolving parasites (due to a short generation time and a large number of offspring) (Frada et al., 2008; Pelusi et al., 2021). Indeed, we observed only two main cellular trajectories of infected cells (Fig. 6): Either, after the rounding of chloroplasts and the appearance of cytosolic RNA spots, we observed bacterial colonies inside the frustule and finally only an empty frustule. Or, a small portion of cells forms auxospores. In the late stages of the infection, we found only few non-auxospore surviving cells.

### 3D biovolumes measurement and implications for biogeochemical cycles

More than a surviving strategy against viral infection, auxospore production also has an important implication for the cycling of organic matter. During viral infection, the cellular content of the infected cells is released into the environment through dissolved organic matter which supplies the pool of organic matter available to prokaryotes and bacterial communities. Our microscopy monitoring shows the development of bacteria that feed on cell debris generated by viral infection. This phenomenon, termed the “viral shunt”, leads to a net increase in bacterial respiration and a decrease in microzooplankton respiration and mitigates the biological pump (Fuhrman, 1999; Wilhelm and Suttle, 1999). Hence, living biomass is diverted away from the higher trophic levels and redirected into the microbial loop. While the hypothesis of the viral shunt has prevailed for many years, several studies suggest that viral infection, including infection of diatoms, can also prime carbon export, this process is termed the viral shuttle, by accelerating sedimentation of the infected host (Lawrence and Suttle, 2004), or via the formation of large particle aggregates or spores (Gobler et al., 1997; Uitz et al., 2010; Laber et al., 2018; Yamada et al., 2018). Considering the importance of *G. delicatula* in temperate coastal ecosystems, determining the relative share of viral shunt versus viral shuttle will be important to understand the fate of this diatom species. The high-throughput 3D measurement of biovolumes of individual cells provides an interesting direct link between single-cell and population-level measurements of virally lysed biomass. Future investigations using this method to compare samples collected from different depths will help to elucidate this question.

### Advantages and limitations of automated confocal microscopy

Understanding the contribution of pathogens to *Guinardia* mortality remains challenging particularly because of the lack of routine methods to detect viral infection. The small size of viruses typically requires Transmission Electron Microscopy (TEM) to reveal their structure. However, these methods offer too little throughput to examine population dynamics. Furthermore, infected diatom cells are very fragile and easily damaged during sample preparation for TEM. 3D multicolour confocal microscopy offers a complementary approach: While it cannot resolve the structure of single viral particles, this technique is suitable to detect viral factories within infected cells as well as three-dimensional morphological changes that occur during infection. Automation of the image acquisition process allows us to image many cells in a relatively short time of analysis (compared to electron microscopy). The detection of chlorophyll autofluorescence, a widely used marker of phytoplankton biomass and water quality, offers an important link between insights into subcellular changes during infection to other observation tools across spatial scales to study key metabolic fluxes. Finally, this method does not require intensive sample processing as fixation and staining procedures are conducted on the sample directly and they are achievable in less than 30 minutes. In this study, we showed that this imaging technique offers great potential for the detection of infected *G. delicatula* or other marine eukaryotic plankton. A remaining limitation of our approach for quantifying virus-induced mortality in the field, is the lack of virus-specific fluorescent labeling, ideally with taxonomic resolution. Future research will therefore focus on its integration with existing methods, such as virus-FISH or qRT-PCR targeting genes of uncultured viruses. Together, these tools should provide new promising high-throughput detection methods to study parasitic infections in environmental samples.

To conclude, viruses are key to (not only marine) ecosystems, from the genetic impact on their hosts to the modification of biogeochemical cycles. They also represent major drivers of biological diversity and evolution and have even been pointed out as a biocontrol agent for emerging infectious diseases (Oliveira et al., 2012; Cohen et al., 2013). They can no longer be ignored in global ecological models. One major limitation to fully understanding their role is the quantification of the mortality that they induce. This innovative study permitted us to expand our understanding of viral infection of the marine diatom *Guinardia delicatula*. The use of high-resolution 3D imaging using confocal microscopy enables the detection of discriminative features of virally infected cells, likely reflecting physiological and metabolic alterations during the infection. Our microscopy monitoring also pointed to the rapid development of escape strategy to viral infection which might have important implications for host population fate.

## Conflict of Interest

The authors declare that the research was conducted in the absence of any commercial or financial relationships that could be construed as a potential conflict of interest.

## Author Contributions

MW, NS and ACB developed the initial idea. NS and ACB prepared algal and viral cultures. MW designed the microscopic imaging experiments and the image analysis. NS and ACB designed the physiological experiments. CC, MW and ACB performed experiments and wrote the initial manuscript. MW performed the morphometric analysis. NS, ACB and CdV provided valuable feedback and criticism on the project development as well as economic resources. All authors revised and contributed to the final version of the manuscript.

## Acknowledgments

Imaging and image analysis were made possible by the MerImage facility at the Station Biologique de Roscoff. We particularly thank Sophie LePanse and Sebastien Colin for the helpful discussions on microscopy and Gilles Mirambeau for the discussions on nucleic acid dyes.

MW was supported by a Benjamin Franklin Fellowship (project 464344344) from the German Research Council (Deutsche Forschungsgemeinschaft, DFG) as well as a *Stratégie d’attractivité durable* (SAD) fellowship from the Région Bretagne.

## Data Availability

All image analysis code is made available under an open-source license (*github.com/mariescopy/Guinardia_ViralInfection*). Data plots shown in this report were created in RStudio. All R scripts and intermediate Excel tables are also available on GitHub. All raw and processed images are publicly available at EBI-BioStudies, accession ID: S-BIAD534. Taxonomically and ecologically annotated projections will also be released on EcoTaxa (*ecotaxa.obs-vlfr.fr*) for public exploration.

## Notes

https://www.ebi.ac.uk/biostudies/studies/S-BIAD534

### Competing Interest Statement

The authors have declared no competing interest.

### Summary of Updates

- We have merged Figure 1 & 4 into one new Figure 1 to make the chronology of events clearer to the readers and updated the Result section about "Cell mortality over time " - A section on "Interplay of bacterial and viral infections" has been added to the Discussion section - Figure 5 (biomass) and associated text have been moved to the Supplementary material

## References

Allers, E., Moraru, C., Duhaime, M. B., Beneze, E., Solonenko, N., Barrero-Canosa, J., et al. (2013). Single-cell and population level viral infection dynamics revealed by phageFISH, a method to visualize intracellular and free viruses. Environ. Microbiol. 15, 2306–2318. doi: 10.1111/1462-2920.12100.

Armbrust, E. V. (2009). The life of diatoms in the world’s oceans. Nature 459, 185–192. doi: 10.1038/nature08057.

Arsenieff, L. (2018). Parasitisme et contrôle des blooms de diatomées en Manche Occidentale.pdf.

Arsenieff, L., Kimura, K., Kranzler, C. F., Baudoux, A.-C., and Thamatrakoln, K. (2022). “Diatom Viruses,” in The Molecular Life of Diatoms, eds. A. Falciatore and T. Mock (Cham: Springer International Publishing), 713–740. doi: 10.1007/978-3-030-92499-7_24.

Arsenieff, L., Simon, N., Rigaut-Jalabert, F., Le Gall, F., Chaffron, S., Corre, E., et al. (2019). First Viruses Infecting the Marine Diatom Guinardia delicatula. Front. Microbiol. 9.

Bar-On, Y. M., and Milo, R. (2019). The Biomass Composition of the Oceans: A Blueprint of Our Blue Planet. Cell 179, 1451–1454. doi: 10.1016/j.cell.2019.11.018.

Caracciolo, M., Rigaut-Jalabert, F., Romac, S., Mahé, F., Forsans, S., Gac, J., et al. (2022). Seasonal dynamics of marine protist communities in tidally mixed coastal waters. Mol. Ecol. 31, 3761–3783. doi: 10.1111/mec.16539.

Cohen, Y., Joseph Pollock, F., Rosenberg, E., and Bourne, D. G. (2013). Phage therapy treatment of the coral pathogen Vibrio coralliilyticus. MicrobiologyOpen 2, 64–74. doi: 10.1002/mbo3.52.

Colin, S., Coelho, L. P., Sunagawa, S., Bowler, C., Karsenti, E., Bork, P., et al. (2017). Quantitative 3D-imaging for cell biology and ecology of environmental microbial eukaryotes. eLife 6, e26066. doi: 10.7554/eLife.26066.

Crawford, R. M. (1995). The role of sex in the sedimentation of a marine diatom bloom. Limnol. Oceanogr. 40, 200–204. doi: 10.4319/lo.1995.40.1.0200.

Frada, M., Probert, I., Allen, M. J., Wilson, W. H., and de Vargas, C. (2008). The “Cheshire Cat” escape strategy of the coccolithophore *Emiliania huxleyi* in response to viral infection. Proc. Natl. Acad. Sci. 105, 15944–15949. doi: 10.1073/pnas.0807707105.

Fuhrman, J. A. (1999). Marine viruses and their biogeochemical and ecological effects. Nature 399, 541–548. doi: 10.1038/21119.

Gobler, C. J., Hutchins, D. A., Fisher, N. S., Cosper, E. M., and Saňudo-Wilhelmy, S. A. (1997). Release and bioavailability of C, N, P Se, and Fe following viral lysis of a marine chrysophyte. Limnol. Oceanogr. 42, 1492–1504. doi: 10.4319/lo.1997.42.7.1492.

Guilloux, L., Rigaut-Jalabert, F., Jouenne, F., Ristori, S., Viprey, M., Not, F., et al. (2013). An annotated checklist of Marine Phytoplankton taxa at the SOMLIT-Astan time series off Roscoff (Western English Channel, France): data collected from 2000 to 2010. Cah. Biol. Mar. 54, 247–256.

Gustavsen, J. A., Winget, D. M., Tian, X., and Suttle, C. A. (2014). High temporal and spatial diversity in marine RNA viruses implies that they have an important role in mortality and structuring plankton communities. Front. Microbiol. 5. doi: 10.3389/fmicb.2014.00703.

Hernández-Fariñas, T., Soudant, D., Barillé, L., Belin, C., Lefebvre, A., and Bacher, C. (2014). Temporal changes in the phytoplankton community along the French coast of the eastern English Channel and the southern Bight of the North Sea. ICES J. Mar. Sci. 71, 821–833. doi: 10.1093/icesjms/fst192.

Julius, M. L., and Theriot, E. C. (2010). “The diatoms: a primer,” in The Diatoms, eds. J. P. Smol and E. F. Stoermer (Cambridge University Press), 8–22. doi: 10.1017/CBO9780511763175.003.

Kimura, K., and Tomaru, Y. (2015). Discovery of Two Novel Viruses Expands the Diversity of Single-Stranded DNA and Single-Stranded RNA Viruses Infecting a Cosmopolitan Marine Diatom. Appl. Environ. Microbiol. 81, 1120–1131. doi: 10.1128/AEM.02380-14.

Kranzler, C. F., Brzezinski, M. A., Cohen, N. R., Lampe, R. H., Maniscalco, M., Till, C. P., et al. (2021). Impaired viral infection and reduced mortality of diatoms in iron-limited oceanic regions. Nat. Geosci. 14, 231–237. doi: 10.1038/s41561-021-00711-6.

Kranzler, C. F., Krause, J. W., Brzezinski, M. A., Edwards, B. R., Biggs, W. P., Maniscalco, M., et al. (2019). Silicon limitation facilitates virus infection and mortality of marine diatoms. Nat. Microbiol. 4, 1790–1797. doi: 10.1038/s41564-019-0502-x.

Kröger, N., and Poulsen, N. (2008). Diatoms—From Cell Wall Biogenesis to Nanotechnology. Annu. Rev. Genet. 42, 83–107. doi: 10.1146/annurev.genet.41.110306.130109.

Laber, C. P., Hunter, J. E., Carvalho, F., Collins, J. R., Hunter, E. J., Schieler, B. M., et al. (2018). Coccolithovirus facilitation of carbon export in the North Atlantic. Nat. Microbiol. 3, 537–547. doi: 10.1038/s41564-018-0128-4.

Lawrence, J., and Suttle, C. (2004). Effect of viral infection on sinking rates of Heterosigma akashiwo and its implications for bloom termination. Aquat. Microb. Ecol. 37, 1–7. doi: 10.3354/ame037001.

Leblanc, K., Arístegui, J., Armand, L., Assmy, P., Beker, B., Bode, A., et al. (2012). A global diatom database – abundance, biovolume and biomass in the world ocean. Earth Syst. Sci. Data 4, 149–165. doi: 10.5194/essd-4-149-2012.

Mann, D. G., and Droop, S. J. M. (1996). “Biodiversity, biogeography and conservation of diatoms,” in Biogeography of Freshwater Algae: Proceedings of the Workshop on Biogeography of Freshwater Algae, held during the Fifth International Phycological Congress, Qingdao, China, June 1994 Developments in Hydrobiology., ed. J. Kristiansen (Dordrecht: Springer Netherlands), 19–32. doi: 10.1007/978-94-017-0908-8_2.

Mayer, K. A., Stöckl, J., Zlabinger, G. J., and Gualdoni, G. A. (2019). Hijacking the Supplies: Metabolism as a Novel Facet of Virus-Host Interaction. Front. Immunol. 10. Available at: https://www.frontiersin.org/articles/10.3389/fimmu.2019.01533 [Accessed November 29, 2022].

Miranda, J. A., Culley, A. I., Schvarcz, C. R., and Steward, G. F. (2016). RNA viruses as major contributors to Antarctic virioplankton. Environ. Microbiol. 18, 3714–3727. doi: 10.1111/1462-2920.13291.

Netherton, C. L., and Wileman, T. (2011). Virus factories, double membrane vesicles and viroplasm generated in animal cells. Curr. Opin. Virol. 1, 381–387. doi: 10.1016/j.coviro.2011.09.008.

Oliveira, J., Castilho, F., Cunha, A., and Pereira, M. J. (2012). Bacteriophage therapy as a bacterial control strategy in aquaculture. Aquac. Int. 20, 879–910. doi: 10.1007/s10499-012-9515-7.

Ollion, J., Cochennec, J., Loll, F., Escudé, C., and Boudier, T. (2013). TANGO: a generic tool for high-throughput 3D image analysis for studying nuclear organization. Bioinformatics 29, 1840–1841. doi: 10.1093/bioinformatics/btt276.

Peacock, E., Olson, R., and Sosik, H. (2014). Parasitic infection of the diatom Guinardia delicatula, a recurrent and ecologically important phenomenon on the New England Shelf. Mar. Ecol. Prog. Ser. 503, 1–10. doi: 10.3354/meps10784.

Pelusi, A., De Luca, P., Manfellotto, F., Thamatrakoln, K., Bidle, K. D., and Montresor, M. (2021). Virus-induced spore formation as a defense mechanism in marine diatoms. New Phytol. 229, 2251–2259. doi: 10.1111/nph.16951.

Reed, L. J., and Muench, H. (1938). A simple method of estimating fifty per cent endpoints. Am. J. Epidemiol. 27, 493–497. doi: 10.1093/oxfordjournals.aje.a118408.

Rohwer, F., and Thurber, R. V. (2009). Viruses manipulate the marine environment. Nature 459, 207–212. doi: 10.1038/nature08060.

Schindelin, J., Arganda-Carreras, I., Frise, E., Kaynig, V., Longair, M., Pietzsch, T., et al. (2012). Fiji: An open-source platform for biological-image analysis. Nat. Methods 9, 676–682. doi: 10.1038/nmeth.2019.

Schlüter, M. H., Kraberg, A., and Wiltshire, K. H. (2012). Long-term changes in the seasonality of selected diatoms related to grazers and environmental conditions. J. Sea Res. 67, 91–97. doi: 10.1016/j.seares.2011.11.001.

Shirai, Y., Tomaru, Y., Takao, Y., Suzuki, H., Nagumo, T., and Nagasaki, K. (2008). Isolation and Characterization of a Single-Stranded RNA Virus Infecting the Marine Planktonic Diatom Chaetoceros tenuissimus Meunier. Appl. Environ. Microbiol. 74, 4022–4027. doi: 10.1128/AEM.00509-08.

Suttle, C. A. (2005). Viruses in the sea. Nature 437, 356–361. doi: 10.1038/nature04160.

Tillmann, U., Hesse, K.-J., and Tillmann, A. (1999). Large-scale parasitic infection of diatoms in the Northfrisian Wadden Sea. J. Sea Res. 42, 255–261. doi: 10.1016/S1385-1101(99)00029-5.

Tinevez, J. Y., Rueden, C., Cardone, G., Schindelin, J., Eglinger, J., and Hiner, M. C. (2018). Directionality. ImageJ Plugin. Available at: https://imagej.github.io/plugins/directionality [Accessed July 14, 2022].

Tomaru, Y., Kimura, K., and Yamaguchi, H. (2014). Temperature alters algicidal activity of DNA and RNA viruses infecting Chaetoceros tenuissimus. Aquat. Microb. Ecol. 73, 171–183. doi: 10.3354/ame01713.

Tomaru, Y., Toyoda, K., and Kimura, K. (2015). Marine diatom viruses and their hosts: Resistance mechanisms and population dynamics. Perspect. Phycol., 69–81. doi: 10.1127/pip/2015/0023.

Tréguer, P., Bowler, C., Moriceau, B., Dutkiewicz, S., Gehlen, M., Aumont, O., et al. (2018). Influence of diatom diversity on the ocean biological carbon pump. Nat. Geosci. 11, 27–37. doi: 10.1038/s41561-017-0028-x.

Uitz, J., Stramski, D., Baudoux, A.-C., Reynolds, R. A., Wright, V. M., Dubranna, J., et al. (2010). Variations in the optical properties of a particle suspension associated with viral infection of marine bacteria. Limnol. Oceanogr. 55, 2317–2330. doi: 10.4319/lo.2010.55.6.2317.

Veldhuis, M., Kraay, G., and Timmermans, K. (2001). Cell death in phytoplankton: correlation between changes in membrane permeability, photosynthetic activity, pigmentation and growth. Eur. J. Phycol. 36, 167–177. doi: 10.1080/09670260110001735318.

Vlok, M., Lang, A. S., and Suttle, C. A. (2019). Marine RNA Virus Quasispecies Are Distributed throughout the Oceans. mSphere 4, e00157–19. doi: 10.1128/mSphereDirect.00157-19.

Warwick-Dugdale, J., Buchholz, H. H., Allen, M. J., and Temperton, B. (2019). Host-hijacking and planktonic piracy: how phages command the microbial high seas. Virol. J. 16, 15. doi: 10.1186/s12985-019-1120-1.

White, J. G., Amos, W. B., and Fordham, M. (1987). An Evaluation of Confocal versus Conventional Imaging of Biological Structures by Fluorescence Light Microscopy. J. Cell Biol. 105, 41–46.

Wilhelm, S. W., and Suttle, C. A. (1999). Viruses and Nutrient Cycles in the Sea. BioScience 49, 781–788. doi: 10.2307/1313569.

Yamada, Y., Tomaru, Y., Fukuda, H., and Nagata, T. (2018). Aggregate Formation During the Viral Lysis of a Marine Diatom. Front. Mar. Sci. 5. doi: 10.3389/fmars.2018.00167.

