## Supplementary Material for "Viral infection impacts the 3D subcellular structure of the abundant marine diatom *Guinardia delicatula*"

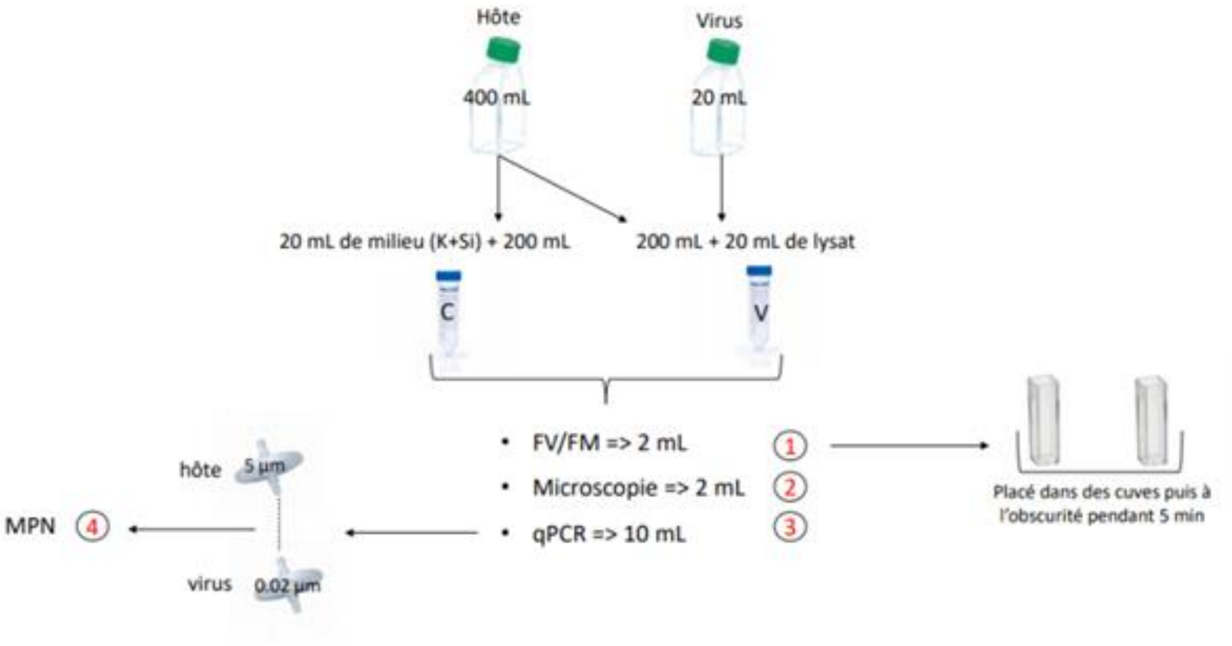


**Supplementary Figure 1**: Overview of viral infection experiments. 400 mL culture of *G. delicatula* in exponential growth was established and divided into two equal aliquots of 200 mL. One aliquot was infected with 20 mL of freshly produced viral lysate. The second served as a control that was amended with 20 mL of K+Si medium. Samples for viral titer, $F_v/F_m$, and microscopy analysis were taken twice daily.

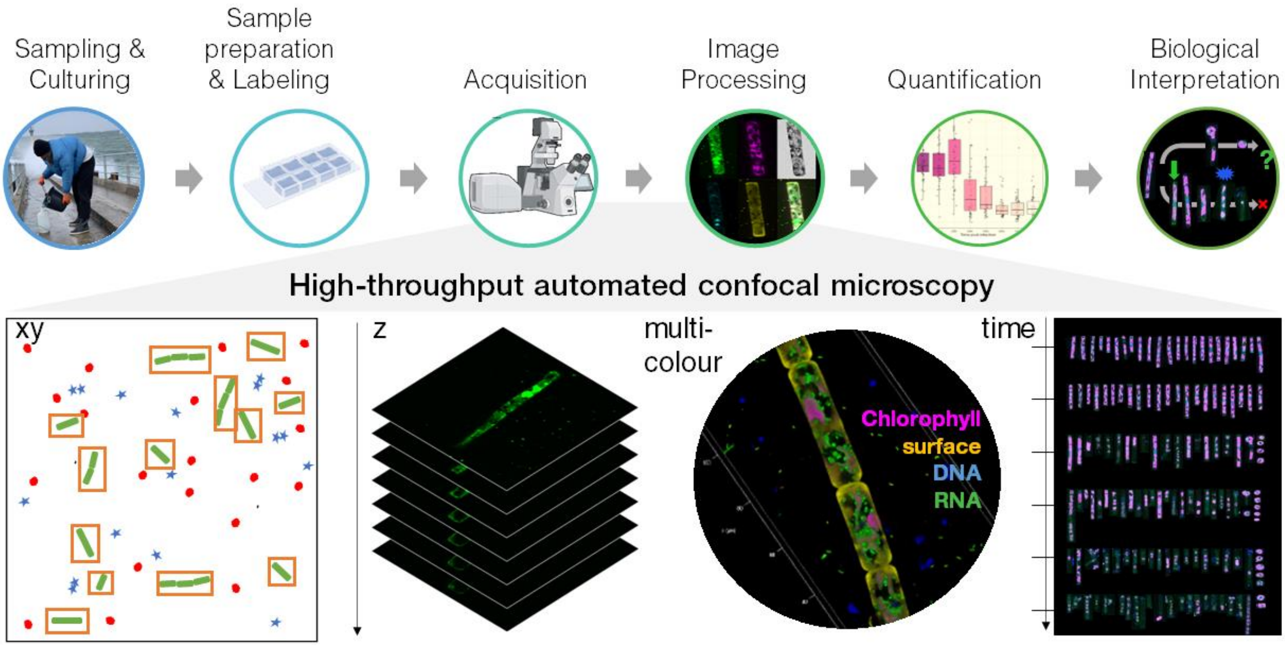


**Supplementary Figure 2: Overview of bioimaging pipeline.** This schematic overview shows the different steps of the imaging approach used in this study.

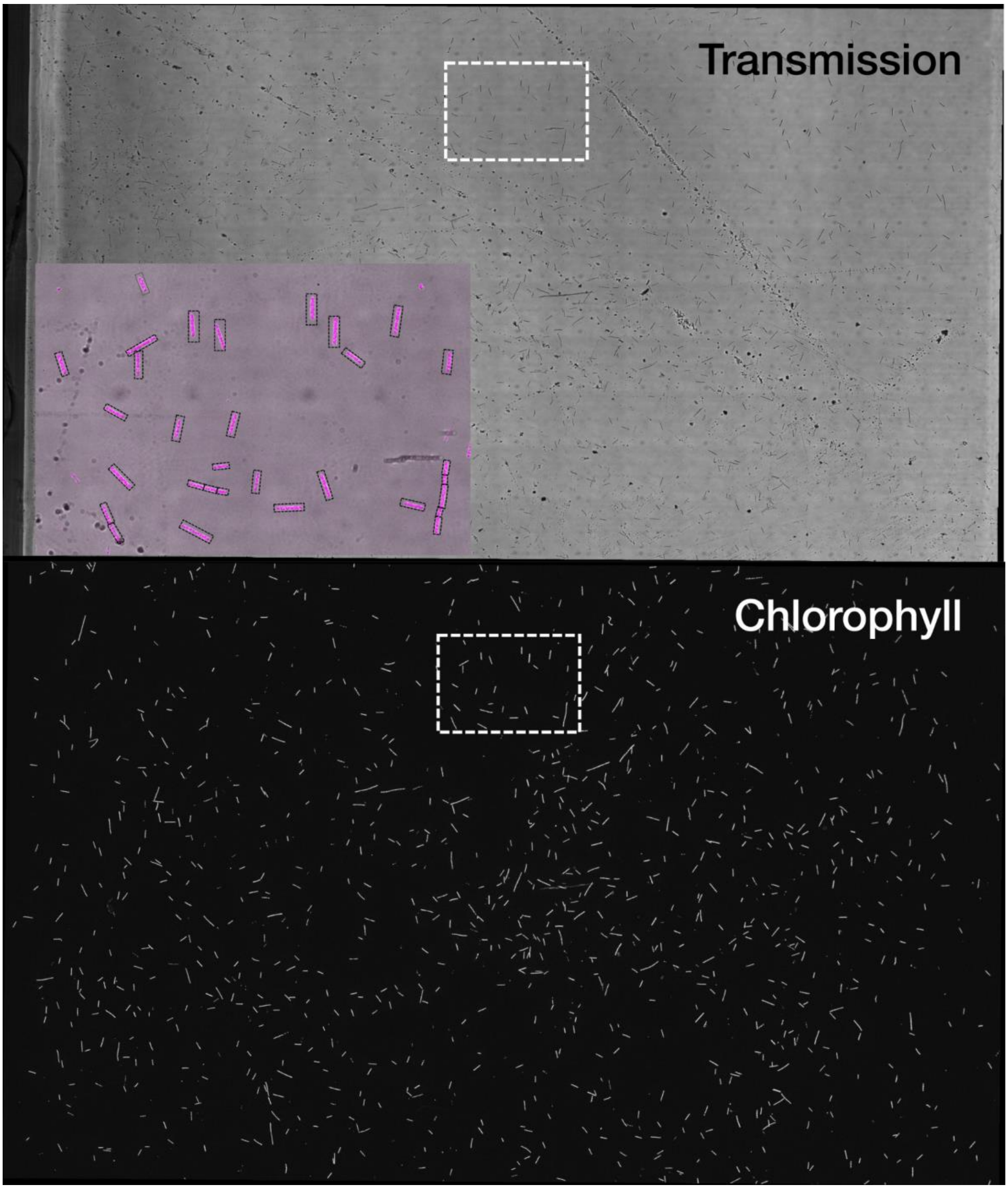


**Supplementary Figure 3: Overview images to estimate abundances.** Cell counts were extracted from low-resolution overview images by automatic particle counting. Here, an overview transmission and Chlorophyll autofluorescence image are shown with a zoom-in overlay inset showing segmented *G. delicatula* cells.

### Supplementary Data 4: Biovolume & biomass estimation

*Guinardia delicatula* cells were approximated as cylindrically shaped as suggested in the global diatom database. *Reproduced from: https://doi.pangaea.de/10.1594/PANGAEA.777384*
*(Creative Commons Licence CC BY 3.0)*

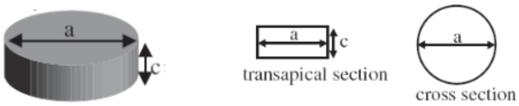


Their diameter and cross-section were measured directly from image data.

### Supplementary Table 4a: Measured cell volumes and chloroplast propostions for each time point

| **Time point** | | **Median cell volume [µm³]** | | **Biomass [ng C / L]** | | **Proportion of chloroplasts %** | |
|---|---|---|---|---|---|---|---|
| | | *Infected* | *Control* | *Infected* | *Control* | *Infected* | *Control* |
| t0 | 0h | 1733 | 1571 | 1.66 | 1.77 | 48 | 42 |
| t1 | 20h | 1617 | 1656 | 1.69 | 1.40 | 49 | 39 |
| t2 | 26h | 1687 | 1420 | 1.76 | 1.84 | 56 | 50 |
| t3 | 44h | 1583 | 1548 | 0.93 | 2.30 | 21 | n.a. |
| t4 | 50h | 1606 | 1930 | 0.50 | 2.53 | 16 | 43 |
| t5 | 68h | 1402 | 1907 | 0.58 | 4.09 | 13 | 45 |
| t6 | 74h | 1431 | 2006 | 0.52 | 4.21 | 12 | 56 |
| t7 | 92h | 1380 | 2193 | 0.28 | 4.03 | 15 | 49 |

### Supplementary Material 5: Biovolumes & biomass plots

Biomass was estimated from measured biovolumes and abundances of diatoms: While the samples from the control culture showed steady growth with a 2.3-fold increase in biomass (1.77 to 4.03 ng C/L) during 92h, the virally infected culture dropped in biomass from initially 1.66 to only 0.52 ng C/L.

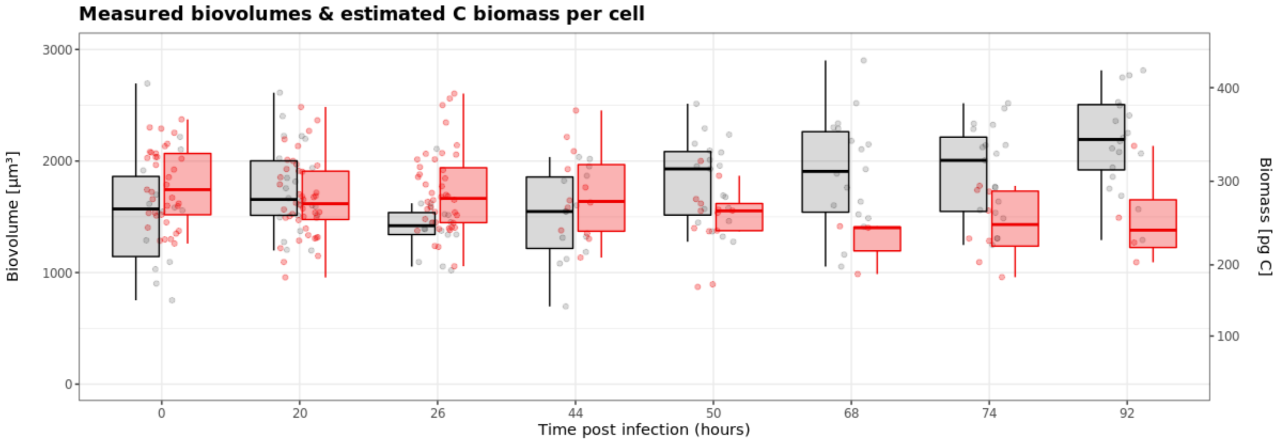


**Supplementary Figure 5a.** Measured biovolumes and estimated C biomass per cell.

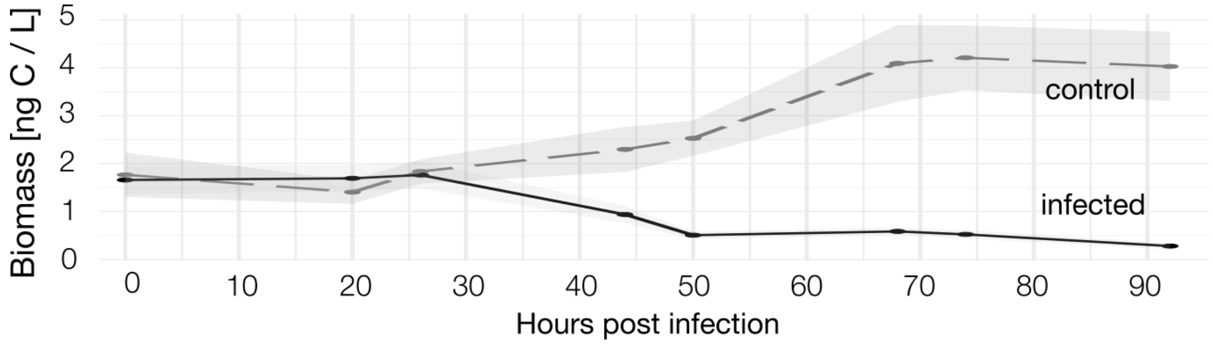


**Supplementary Figure 5b**: Total biomass lysed in an infected cultured compared to control samples

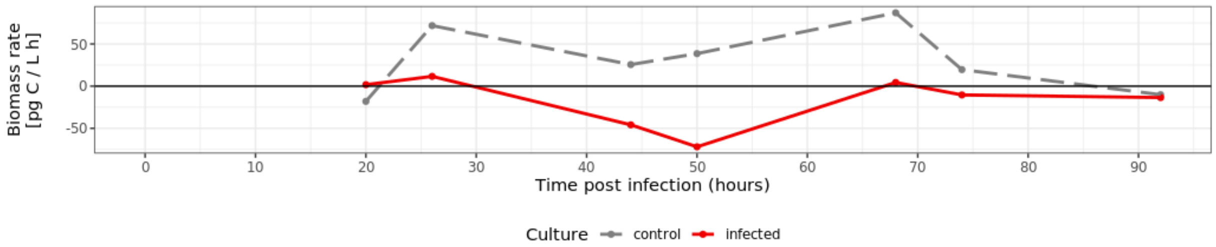


**Supplementary Figure 5c.** Estimated rate of biomass production during the course of the infection.

**Supplemetary Material 6: RNA dyes**

We tested a series of nucleic acid dyes with very similar spectral properties (SybrGreen, PicoGreen, StrandBrite and SYTO RNASelect) regarding

(i) their compatibility with formaldehyde fixation and

(ii) their specificity for ssRNA inside infected and non-infected diatoms in comparison to a DNA stain (Hoechst33342).

(iii) Signal strength over the chlorophyll autofluorescence background

The fluorescence spectra of compared RNA dyes all share very similar parameters, (maximum excitation around 490 nm; emission peak 520-530 nm) which enabled a direct comparison of their suitability to label accumulations of single-stranded virus RNA accumulations inside of infected diatoms. We found that all dyes fulfilled criterion (i) and showed similar results for (ii). The most critical factor was criterion (iii). In our hands and for our biological system, we achieved the best signal-to-noise ratio with StrandBrite.

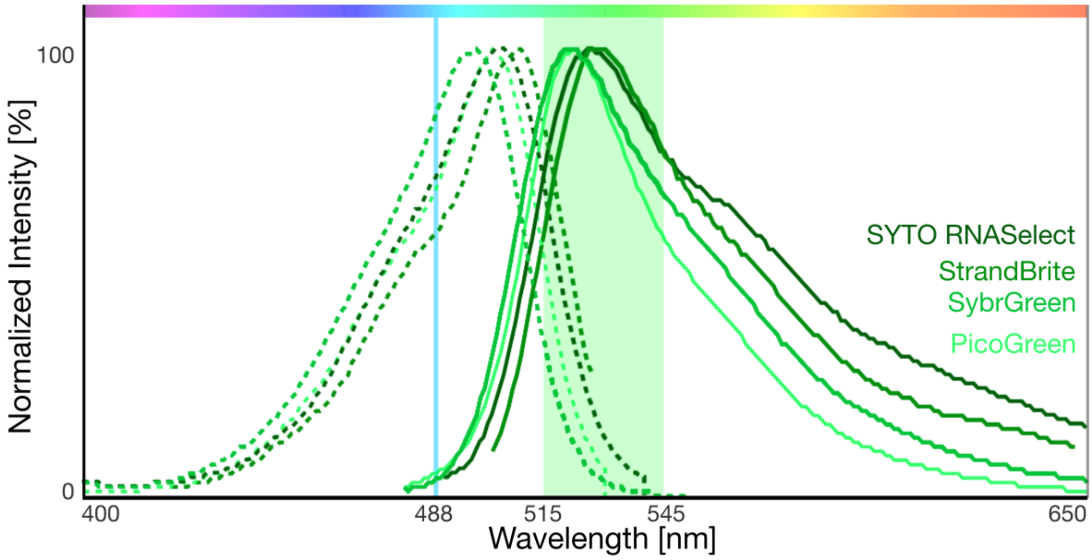
